# Thermal Similarity Attenuates the Strength of Bergmann’s Rule in Birds

**DOI:** 10.64898/2026.11.12.766076

**Authors:** Shanshan Yang, Tariq Ahmad, Haiqin Qin, Zihan Jiang

**Affiliations:** CAS Key Laboratory for Plant Diversity and Biogeography of East Asia; Kunming Institute of Botany, Chinese Academy of Sciences, Kunming, China; Yunnan Research Innovation Promotion Program for State Key Laboratory of Plant Diversity and Specialty Crops; College of Wildlife and Protected Area, Northeast Forestry University, Harbin 150040, China

**Author notes:** Corresponding author: Zihan Jiang.

**Keywords:** body mass, temperature gradient, ecological divergence, congeneric species, macroecology, ecogeographical rule

## Abstract

Geographic body-size variation, classically framed as Bergmann’s rule, is a fundamental indicator of temperature-mediated morphological adaptation. However, empirical support for Bergmann’s rule in birds remains highly unpredictable across lineages. We hypothesized that this variation is caused by differences in thermal similarity among species within the same genus, which may limit temperature-driven changes in body size and be influenced by ecological factors. Using a global dataset comprising 5,715 bird species across 513 genera, we observed that thermal similarity was the strongest predictor of variation in the strength of Bergmann’s rule among competing hypotheses. Among genera exhibiting significantly negative body mass–temperature relationships, higher thermal similarity was consistently associated with less negative slopes, indicating weaker Bergmannian clines. Subgroup analyses indicated that the positive thermal similarity–slope association was supported in dense and semi-open habitats and among non-migratory birds, but was not clearly supported in open habitats or among migratory birds. Our findings provide a novel explanation for variation in the strength of Bergmannian clines among avian lineages, showing that stronger morphological differentiation along thermal gradients is associated with greater thermal contrast. Consequently, our findings provide a basis for testing whether incorporating thermal similarity and ecological context can improve forecasts of avian morphological responses to ongoing climate change.

**Significance Statement:** This study addresses a central question in spatial macroecology: why Bergmannian body size–temperature clines vary in strength among closely related bird lineages. Using a global dataset of 5,715 species in 513 genera and a two-stage comparative framework, we show that, among genera exhibiting significantly negative clines, greater thermal similarity is associated with less negative body mass–temperature slopes, indicating weaker clinal differentiation. This association is most clearly supported in dense and semi-open habitats and among non-migratory genera. By extending the global assessment of He et al. (2023) from the broad occurrence of body size–climate relationships to variation in their strength, the study identifies variation in thermal similarity as a potential source of heterogeneity in a classic spatial ecological pattern. Our findings highlight how within-lineage ecological structure shapes broad geographical patterns. This mechanistic focus on variation in spatial ecogeographical relationships makes the study directly relevant to *Ecography*.

## Introduction

An animal’s body size shapes nearly every aspect of its physiology, ecology, and life history, underlying much of the interspecific variation in metabolism, energetics, and population dynamics (Kozłowski et al. 2020, Weil et al. 2023, White & Marshall 2023). More than 150 years ago, Bergmann (1848) observed that within closely related groups of endotherms, species inhabiting colder regions tend to be larger than their relatives in warmer regions. Birds provide a particularly compelling system for testing the mechanisms behind this rule. As endotherms with high metabolic demands, birds theoretically face strong temperature-mediated selection for heat conservation (Salewski & Watt 2017). However, their exceptional mobility allows them to exploit a wide range of habitat structures and migratory strategies, enabling them to decouple ecologically and behaviorally from macroclimatic extremes (Mainwaring & Street 2021, Rodrigues & Botero 2025). Consequently, empirical support for the rule in avian clades remains highly inconsistent. Rather than a universal pattern, the strength of Bergmannian clines varies drastically across avian lineages, leaving its general applicability heavily contested (Riemer et al. 2018, He et al. 2023, Probst et al. 2026).

To date, multiple factors have been proposed to explain this pervasive heterogeneity in clinal strength, generally falling into two overarching categories. First, conventional explanations typically focus on external environmental variables, suggesting that spatial variation in energy availability, precipitation, and local topography can mask temperature-mediated selection (Meiri & Dayan 2003, Riemer et al. 2018, He et al. 2023). Second, macroevolutionary perspectives emphasize intrinsic evolutionary attributes. Pervasive phylogenetic constraints, limited phenotypic evolutionary rates, or a young clade age can inherently restrict a lineage’s evolutionary capacity to respond to climatic gradients. Under this temporal framework, a recently diverged lineage or one subject to strong morphological inertia may simply lack sufficient evolutionary flexibility or time to accumulate distinct body-size adaptations along contemporary thermal gradients (Gohli & Voje 2016, Pienaar et al. 2026).

However, these explanations often overlook a basic spatial prerequisite. Ambient temperature can drive morphological divergence only when related species occupy different thermal environments. Macroecological studies frequently assume that a broad geographic distribution automatically exposes a lineage to diverse thermal regimes. Yet, geographic distance does not guarantee thermal contrast. Because closely related species often share baseline physiological tolerances (Saban et al. 2023), an avian genus can span a large continental gradient without its species necessarily partitioning that thermal space. In practice, the temperature distributions that congeneric species experience across their ranges can overlap substantially (Srinivasan et al. 2018). If related species experience similar thermal conditions, the divergent selective pressure needed to drive body-size evolution is simply absent—regardless of range size or other environmental pressures (White & Butlin 2021).

Evaluating the potential for temperature-mediated selection therefore requires quantifying the proportional overlap between the actual temperature distributions of congeneric species, which we define as thermal similarity. We hypothesize that thermal similarity predicts the strength of Bergmannian divergence. In genera with high thermal similarity, overlapping temperature exposures weaken divergent thermal selection. This shared climatic space allows other ecological drivers, such as local resource availability, to shape body size, flattening the mass–temperature slope. Conversely, species in genera with low thermal similarity partition their geographic distributions across distinct temperature regimes. This thermal contrast generates the directional selection required to yield the steeper negative slopes predicted by Bergmann’s rule (Figure 1).

**Figure 1.**
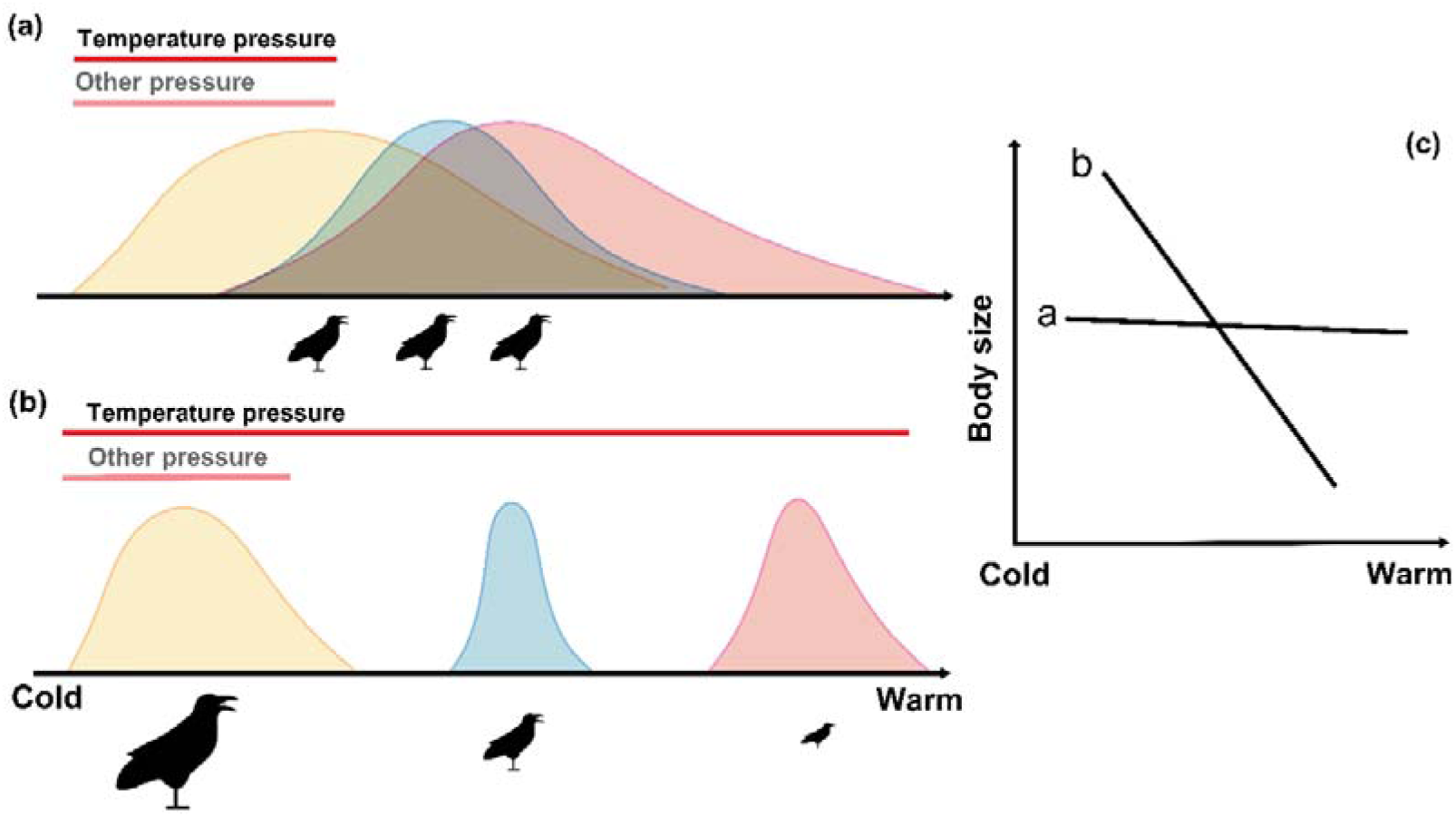
Conceptual framework linking thermal similarity to the strength of Bergmann’s rule in congeneric birds. Three hypothetical congeneric species distributed across a macroclimatic temperature gradient. Colored density curves depict species-specific thermal conditions, with their shaded overlap conceptually defining thermal similarity. In scenario (a), high thermal similarity indicates that species share largely analogous thermal environments. This shared residency minimizes divergent thermal selection, restricting the evolutionary scope for spatial body-size divergence. Conversely, low thermal similarity (b) reflects pronounced macroclimatic segregation. This environmental partitioning drives strong divergent selection, triggering marked morphological adaptation (i.e., larger species evolving in colder regimes and smaller species in warmer ones). (c) The translation of these evolutionary dynamics into expected macroecological clines. High thermal similarity yields a blunted, shallow body size–temperature slope (line a), whereas low thermal similarity facilitates the steep, negative slope classically predicted by Bergmann’s rule (line b).

The predictive strength of thermal similarity, however, likely depends on ecological context. Habitat structure alters how large-scale ambient air temperatures translate into the operative temperatures animals actually experience. In open habitats, animals face direct solar radiation and unpredictable microclimatic fluctuations (Oboikovitz & Swanson 2022). This local thermal noise can decouple operative temperatures from macroclimatic grids, potentially obscuring the macroecological link between thermal similarity and Bergmann’s rule. Closed-canopy forests, by contrast, act as environmental filters that buffer these transient extremes (Frey et al. 2016, De Frenne et al. 2019). By filtering microclimatic noise, stable environments allow geographic temperature gradients to exert a clearer selective pressure on body size. Behavioral strategies similarly uncouple species from local climates. Migration allows birds to track favorable conditions and avoid severe winter exposure (Culp et al. 2017), relaxing temperature-mediated selection. We therefore predict that the positive association between thermal similarity and clinal strength will be most evident in resident lineages and those occupying dense habitats, but weak or absent in migratory species and open habitats.

Here, we analyzed a global dataset of 5,715 bird species across 513 genera to evaluate whether thermal similarity is associated with variation in the strength of Bergmann’s rule among avian genera exhibiting significantly negative body mass–temperature relationships. We quantified genus-level body mass–temperature slopes and pairwise thermal similarity among congeneric species and evaluated their association using Bayesian phylogenetic models that accounted for shared evolutionary history and uncertainty in genus-level slope estimates. Specifically, we addressed three questions: (1) Is thermal similarity associated with variation in the strength of Bergmannian body mass–temperature clines across avian genera? (2) Does thermal similarity receive greater predictive support than alternative environmental and ecological predictors of clinal strength? and (3) Is the thermal similarity–slope association consistently supported across ecological contexts, specifically habitat-openness and migratory-status subgroups? We further assessed the robustness of the primary association using alternative thermal similarity metrics, minimum species-per-genus thresholds, body-mass treatments, and an analysis including all eligible genera irrespective of slope direction or statistical significance.

## Materials and Methods

### Data sources and taxonomic standardization

We assembled a comprehensive global dataset encompassing species-level functional traits, geographic ranges, environmental conditions, and phylogenetic history. Species-level body mass, ecological traits (habitat openness, migratory status, dietary guild, and climate zone), and geographic-range polygons were obtained from the global avian dataset compiled by He et al. (2023). For environmental data, we obtained macroclimatic variables (mean annual temperature, temperature seasonality, and mean annual precipitation) from WorldClim v2.1 at a 30-arc-second resolution (Fick & Hijmans 2017), and net primary productivity (NPP) from the 500-m MODIS/Terra product (Running & Zhao 2021). Avian phylogenies were sourced from the posterior distribution of species-level trees based on the Hackett backbone via BirdTree (Hackett et al. 2008, Jetz et al. 2012). Environmental raster extraction and range-level summary statistics were performed in Python version 3.13.13 using GeoPandas, Rasterio, NumPy, and pandas (McKinney 2010, Harris et al. 2020, Gillies 2023, Jordahl et al. 2024, The pandas development team 2025).

To ensure analytical consistency across morphological, spatial, and phylogenetic datasets, we implemented a highly rigorous taxonomic standardization procedure. Ecological and spatial datasets were first standardized to the IUCN Red List (Version 2024-2) backbone. Subsequently, to link these data with evolutionary history, historical BirdTree taxonomy was reconciled using exact binomial matching and explicit taxonomic crosswalks against BirdLife taxonomic v3 and v10 (http://www.birdlife.org), the IOC World Bird List v15.2 (https://www.worldbirdnames.org/new/), and AviList v2025b (https://www.avilist.org/checklist/v2025b/). We retained only unambiguous one-to-one species correspondences. Any cases involving unresolved splits, lumps, conflicting classifications, or multiple candidate matches were completely excluded to avoid assigning incorrect ecological attributes or phylogenetic positions.

Finally, we aggregated the data to the genus level because the original interspecific formulation of Bergmann’s rule focused on congeneric comparisons. Gridded climatic layers generally fail to adequately capture the operative thermal environments of marine and pelagic birds (Assis et al. 2018, Cox et al. 2018); thus, these species were removed entirely. We restricted our analyses to genera containing at least five valid species, yielding a final core dataset of 5,715 species distributed across 513 genera for all downstream analyses.

### Quantifying the strength of Bergmann’s rule and thermal similarity

To quantify the strength of Bergmann’s rule, we first calculated the median annual temperature across the geographic range of each species. The median reduced the influence of localized temperature extremes within large geographic ranges and was therefore preferred over the mean. We then fitted genus-specific ordinary least-squares regressions of log_10_-transformed body mass against median annual temperature. The resulting slope (β) provided a continuous measure of clinal strength (Mayr 1963, Meiri & Dayan 2003). We classified a genus as conforming to Bergmann’s rule when its slope was negative, and the corresponding genus-specific OLS regression was statistically significant (β < 0 and *p* < 0.05). For the subsequent thermal similarity–slope analyses, both the slope estimate and its standard error were carried forward into the Bayesian phylogenetic models.

To quantify thermal similarity, we converted the grid-cell temperatures within each species’ range into relative frequency distributions following Broennimann et al. (2012). Temperature distributions comprised 1 °C bins, with each bin expressed as the proportion of grid cells *p_i_ = n_i_* / *N* and empty bins assigned a value of zero. Schoener’s *D* quantified pairwise thermal similarity between congeneric species, ranging from zero for no overlap to one for complete overlap (Schoener 1968, Warren et al. 2008). The mean of all pairwise values provided a single thermal similarity estimate for each genus. The geographic distribution and ecological composition of the retained genera are summarized in Figure 2a, b, showing their global distribution, thermal similarity values, and the representation of Bergmannian and non-Bergmannian genera across habitat and migratory-status categories.

**Figure 2.**
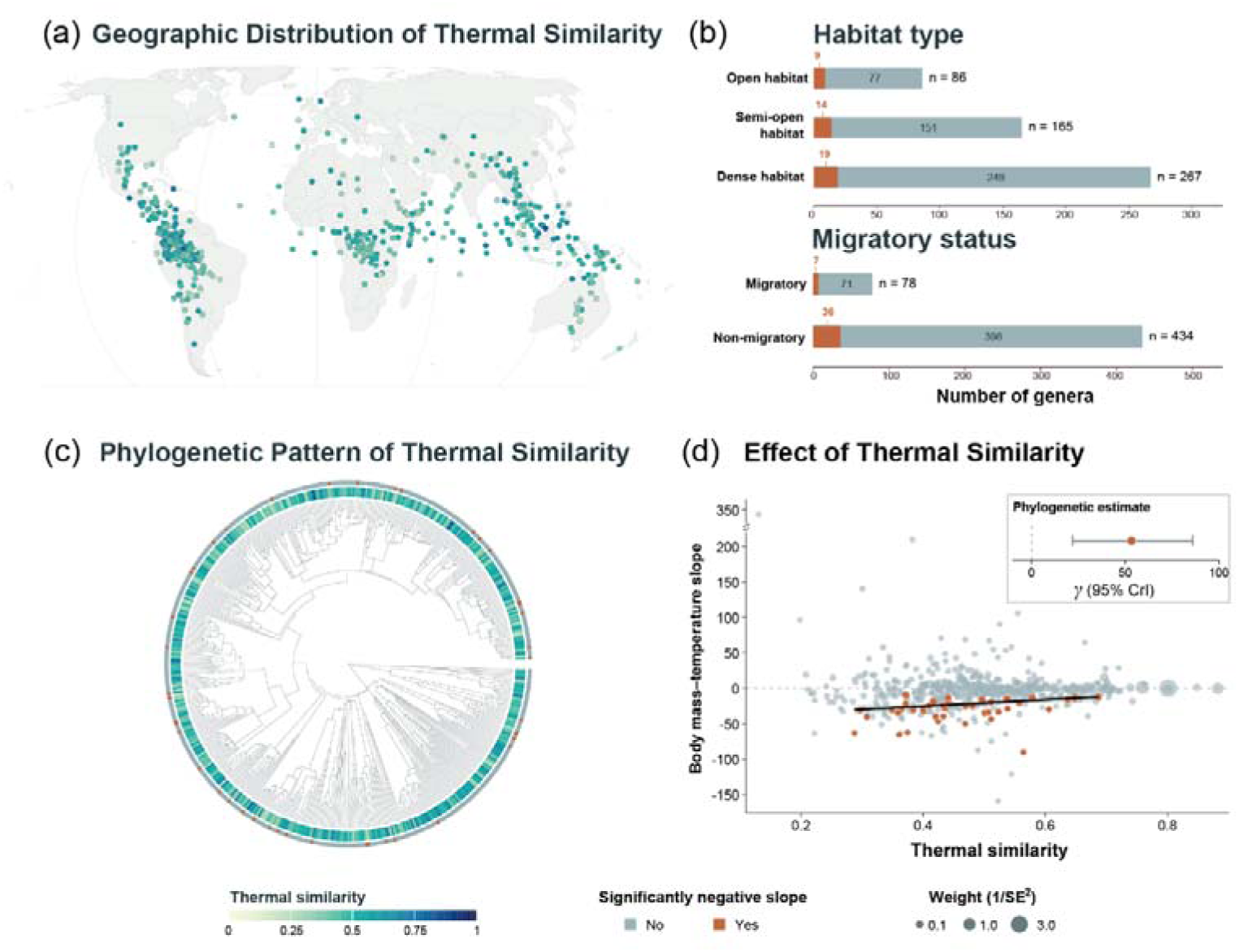
Global distribution, ecological composition, phylogenetic pattern, and association of thermal similarity with Bergmannian clinal strength in birds. (a) Geographic distribution of retained genera, with point color indicating genus-level mean thermal similarity. (b) Numbers of genera with and without significantly negative body mass–temperature slopes (β < 0, *p* < 0.05) across habitat types and migratory categories. (c) Phylogenetic distribution of genus-level thermal similarity and significantly negative slopes. (d) Relationship between thermal similarity and genus-level body mass–temperature slopes; point size is proportional to inverse slope variance (1/SE²), and the black line shows the weighted regression among genera with significantly negative slopes. The inset shows the pooled Bayesian phylogenetic estimate of the thermal similarity coefficient and its 95% credible interval.

### Phylogenetic framework and thermal similarity–slope analysis

To account for non-independence arising from shared ancestry, all downstream inferential models were fitted as Bayesian phylogenetic models. Avian phylogenies came from the posterior distribution of species-level trees based on the Hackett backbone and available through BirdTree (Hackett et al. 2008, Jetz et al. 2012). Because the analyses were conducted at the genus level while the phylogenies were resolved at the species level, one representative species was randomly selected from each retained genus. We sampled 100 of the 1,000 posterior trees without replacement and repeated the representative-species selection ten times per tree, yielding 1,000 phylogenetic realizations and propagating uncertainty from both phylogenetic-tree choice and genus representation. Each analysis retained only genera that could be matched across all 100 sampled trees, thereby maintaining a constant taxonomic sample among realizations. For each realization, the selected tree was pruned to the representative species, tip labels were replaced by genus names, and a genus-level phylogenetic variance–covariance matrix based on shared branch lengths was calculated and scaled to a mean diagonal value of one. Each model included a genus-level random effect structured by this covariance matrix. Phylogenetic processing and Bayesian model fitting were conducted in R (version 4.5.2) using the ape and brms packages, with Rstan (version 2.32.7) as the Stan backend (Bürkner 2017, Carpenter et al. 2017, Paradis & Schliep 2019, R Core Team 2025, Stan Development Team 2025).

Genera exhibiting significantly negative body mass–temperature slopes (β < 0 and p < 0.05) were analyzed using Bayesian phylogenetic meta-regression to test whether mean-centered thermal similarity explained variation in clinal strength (Adams 2008, Hadfield & Nakagawa 2010). The response was specified in brms as Slope | se(Slope_SE, sigma = TRUE), thereby incorporating uncertainty in the genus-level slope estimates while allowing additional residual heterogeneity. Zero-centered normal priors were specified for the intercept and thermal similarity coefficient (γ), and Student-t priors with three degrees of freedom were assigned to the phylogenetic and residual standard deviations (Gelman 2006); the data-scaled prior definitions are provided in the Supporting Information.

Models were fitted with four Markov chains and 4,000 iterations per chain and convergence was evaluated using R, effective sample sizes, and divergent transitions (Vehtari et al. 2021). To summarize uncertainty across phylogenetic realizations, 200 posterior draws were sampled from each converged model, from which posterior means and 95% credible intervals were calculated (de Villemereuil et al. 2012). Convergence diagnostics and posterior-draw processing were implemented using the posterior package (Bürkner et al. 2026).

### Comparison with alternative predictors

To address whether thermal similarity provided greater predictive support than alternative hypotheses, we constructed five candidate models using the previously acquired environmental and trait data: mean annual precipitation, NPP, temperature seasonality, climate zone, and dietary guild. For the continuous climatic variables (precipitation and temperature seasonality), we calculated the median raster value across each species’ geographic range. For NPP, we calculated the mean value across the range to better reflect cumulative resource availability. We then averaged these species-level values to obtain genus-level continuous predictors. We directly aggregated the categorical predictors (dietary guild and climate zone) to the genus level based on the foundational trait classifications. Genera lacking unambiguous classifications were excluded from this specific complete-case model comparison.

All candidate continuous predictors were z-standardized. Separate candidate models were fitted to the identical complete-case set of genera exhibiting significantly negative slopes, using the exact same phylogenetic covariance matrices and Slope | se(Slope_SE, sigma = TRUE) response specification. Pareto-smoothed importance-sampling leave-one-out cross-validation (PSIS-LOO) quantified relative predictive performance. Within each phylogenetic realization, ΔLOOIC and its standard error were calculated relative to the best-supported model, and stacking weights quantified the relative predictive support across the candidate ensemble. PSIS-LOO and stacking calculations were implemented using the loo package (Vehtari et al. 2017, Yao et al. 2018).

### Ecological subgroup analyses

To address the modifying role of ecological contexts, we partitioned the dataset according to the pre-defined habitat openness (dense, semi-open, open) and migratory status (migratory, non-migratory). Habitat-openness analyses were conducted by retaining only species assigned to the corresponding habitat category and recalculating genus-level body mass–temperature slopes and mean thermal similarity within each habitat-specific subset. Genera represented by fewer than five species within a given habitat category were excluded. Because species-level partitioning by migratory status left too few qualifying migratory genera for model estimation (*n* = 2), migratory status was instead assigned at the genus level using a majority rule, with genera containing >50% migratory species classified as migratory and all others as non-migratory. All eligible species within each genus were retained when calculating genus-level slopes and thermal similarity, and genera containing fewer than five species were excluded. The identical Bayesian phylogenetic meta-regression models defined in the primary analysis were then fitted separately to each qualifying subgroup.

### Sensitivity analyses

We conducted four sets of sensitivity analyses to verify the robustness of the primary thermal similarity–slope association. First, we recalculated pairwise thermal similarity using the corrected modified Hellinger distance, Horn’s *R*, and Pianka’s *O*, as defined in Supporting information. Second, we tested minimum thresholds of four and six species per genus to assess sensitivity to our primary five-species requirement. Third, we excluded species with body-mass values derived from morphological correlates or phylogenetic imputation, repeating the analyses strictly using directly observed values or literature-derived values. Fourth, to confirm that our findings were not exclusively driven by truncating the dataset to lineages with significantly negative clines, we repeated the overarching Bayesian phylogenetic analysis across all eligible genera simultaneously, irrespective of slope direction or initial statistical significance. All sensitivity models followed identical MCMC sampling, convergence criteria, and phylogenetic error propagation procedures.

## Results

### Thermal similarity–slope association

The dataset comprised 5,715 bird species assigned to 513 genera distributed globally (Figure 2a). Genus-level body mass–temperature slopes were available for 512 genera, of which 310 had negative slopes and 202 had positive slopes. Of these genera, 43 had negative slopes with *p* < 0.05, 16 had positive slopes with *p* < 0.05, and 453 had *p* ≥ 0.05. The 43 genera with negative slopes and *p* < 0.05 were retained in the primary analysis. Within this primary subset, thermal similarity ranged from 0.288 to 0.687, with a median of 0.441. In the primary Bayesian phylogenetic meta-regression, the thermal similarity coefficient was positive (γ = 53.34, 95% CrI = [21.91, 86.04], Pr[γ > 0] > 0.999; Figure 2d), and the mean Bayesian *R*² across phylogenetic realizations was 0.138.

### Comparison with alternative predictors

Among the six candidate models, the thermal similarity model had the lowest mean LOOIC and the largest mean stacking weight (Table 1). Its mean LOOIC was 339, its mean ΔLOOIC was 0.00, and its mean stacking weight was 0.600. The mean-annual-precipitation model had a mean LOOIC of 342, a mean ΔLOOIC of 3.27, and a mean stacking weight of 0.388. The dietary-guild model also had a mean LOOIC of 342, with a mean ΔLOOIC of 3.14 and a mean stacking weight of 0.012. The climate-zone and temperature-seasonality models had mean LOOIC values of 345 and 346 and mean ΔLOOIC values of 5.95 and 6.84, respectively. The net-primary-productivity model had the highest mean LOOIC (348) and a mean ΔLOOIC of 8.61; the mean stacking weights of these three models were all below 0.001.

**Table 1.** Comparison of phylogenetic models evaluating candidate predictors of variation in genus-level body mass–temperature slopes among genera exhibiting significantly negative body mass–temperature relationships (β < 0, *p* < 0.05).

| Candidate predictor | Mean LOOIC | ΔLOOIC | Stacking weight | Hypothesis literature |
|---|---|---|---|---|
| **Thermal Similarity** | **339** | **0.00** | **0.600** | **——** |
| Dietary Guild | 342 | 3.14 | 0.012 | (He et al. 2023) |
| Mean Annual Precipitation | 342 | 3.27 | 0.388 | (Watson & Kerr 2025) |
| Climate Zone | 345 | 5.95 | < 0.001 | (He et al. 2023) |
| Temperature Seasonality | 346 | 6.84 | < 0.001 | (Skinner et al. 2025) |
| Net Primary Productivity | 348 | 8.61 | < 0.001 | (Henry et al. 2023) |

Each candidate predictor was evaluated in a separate phylogenetic model using the same genus-level slope response and uncertainty structure. Predictive performance was assessed with PSIS-LOO across phylogenetic realizations. Mean LOOIC summarizes average predictive fit, ΔLOOIC gives the difference from the best-supported model, and stacking weights indicate relative predictive support. Boldface denotes the best-supported model; hypothesis literature lists representative studies supporting each predictor.

### Ecological subgroup analyses

Thermal similarity coefficients were estimated separately for three habitat categories and two migratory-status categories (Figure 3). In dense habitats, the thermal similarity coefficient was γ = 72.41 (95% CrI = [22.66, 123.49], *n* = 19), and the mean Bayesian *R*² was 0.391. In semi-open habitats, the coefficient was γ = 144.42 (95% CrI = [44.62, 245.54], *n* = 14), and the mean Bayesian *R*² was 0.676. The 95% CrIs for both habitat categories excluded zero. In open habitats, the coefficient was γ = 34.92 (95% CrI = [-44.93, 108.55], *n* = 9), and the mean Bayesian *R*² was 0.264. Among non-migratory genera, the thermal similarity coefficient was γ = 59.66 (95% CrI = [25.06, 95.74], *n* = 36), and the mean Bayesian *R*² was 0.295. Among migratory genera, the coefficient was γ = 14.50 (95% CrI = [−248.08, 252.49], *n* = 7), and the mean Bayesian *R*² was 0.377. The migratory estimate had the widest 95% CrI among the five subgroup estimates.

**Figure 3.**
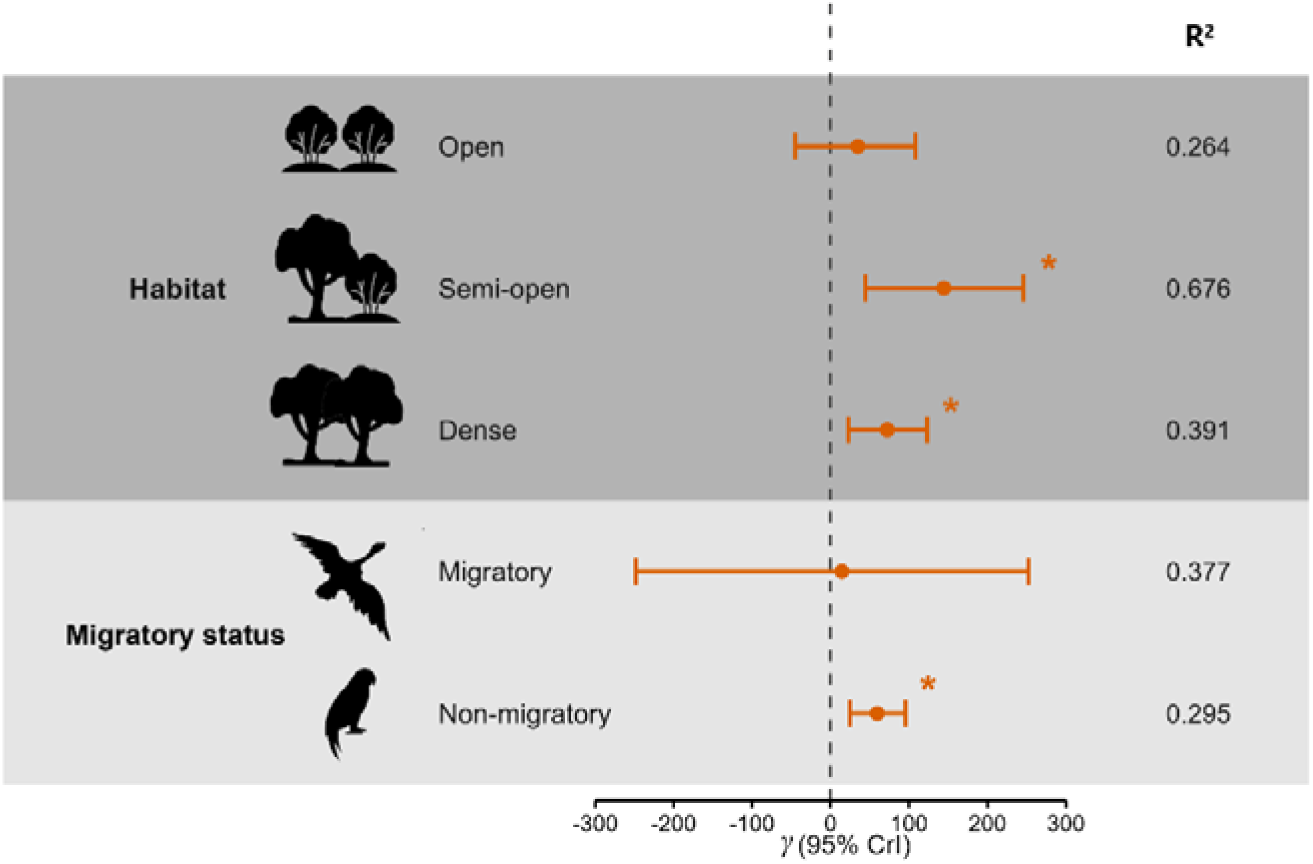
Subgroup-specific effects of thermal similarity on Bergmannian clinal strength in birds. Points and horizontal bars show posterior estimates of the thermal similarity coefficient (γ) and 95% credible intervals (CrIs), respectively, for habitat-openness and migratory-status subgroups. Models included genera with significantly negative body mass–temperature slopes (β < 0, *p* < 0.05), incorporated slope-estimation uncertainty and phylogenetic covariance, and propagated uncertainty across 100 posterior trees and 10 representative-species selections per tree. The dashed line denotes γ = 0; asterisks indicate 95% CrIs excluding zero. Values on the right are mean Bayesian *R*².

### Sensitivity analyses

The principal thermal similarity–slope association was broadly consistent across the retained Bayesian phylogenetic sensitivity analyses. Recalculating thermal similarity using the corrected modified Hellinger distance (γ = 57.32, 95% CrI = [24.34, 91.48]), Horn’s *R* (γ = 46.79, 95% CrI = [19.76, 74.76]), and Pianka’s *O* (γ = 45.06, 95% CrI = [18.09, 72.86]) produced coefficients with the same direction as the primary Schoener’s *D* analysis. With a minimum threshold of six species per genus, the thermal similarity coefficient was γ = 60.32 (95% CrI = [30.17, 92.25]). When the threshold was reduced to four species, the estimated effect remained positive but became more uncertain, with its 95% credible interval including zero (γ = 50.08, 95% CrI = [−9.65, 108.22]). After species with imputed body-mass values were excluded, the posterior thermal similarity coefficient remained positive (γ = 61.01, 95% CrI = [33.61, 88.84]), indicating that the primary result was not driven by body-mass imputation. In addition, in the analysis including all 512 genera with available slope estimates the coefficient was γ = 12.86 (95% CrI = [2.23, 23.59], Pr[γ > 0] = 0.991), and the mean Bayesian *R*² was 0.005. Full results of the sensitivity analyses are provided in the Supporting information.

## Discussion

Our global macroecological analysis helps resolve a persistent ambiguity in the application of Bergmann’s rule: why do avian genera exhibiting significant negative body mass–temperature relationships differ so markedly in the strength of their clines? Among these genera, higher thermal similarity was associated with less negative body mass–temperature slopes, indicating weaker clinal differentiation in body mass. These results suggest that thermal similarity is closely associated with the magnitude of morphological divergence along thermal gradients.

One possible explanation for this pattern is that thermal similarity measures the amount of thermal variation available for temperature-mediated divergence among congeneric species (White & Butlin 2021). Genera with high thermal similarity contain species whose geographic distributions overlap substantially in thermal space, reducing effective thermal contrast across species even when the genus as a whole spans a broad geographic temperature gradient (Salewski & Watt 2017). Under such conditions, changes in ambient temperature among species may provide comparatively weak divergent selection on body size, permitting other environmental factors to contribute more strongly to morphological differentiation. Lower thermal similarity, in contrast, implies greater thermal differentiation among congeneric species, which may provide a stronger ecological setting for temperature-associated body-size divergence. Consequently, the strength of a Bergmannian cline may depend on both the total temperature range occupied by a genus and the degree to which its component species differ in the thermal environments they inhabit. This explanation is consistent with the comprehensive view that ecological differences among closely associated species can mediate the strength of morphological divergence and ecological selection (Sottas et al. 2018).

A complementary justification is that greater thermal similarity may coincide with greater geographic overlap among closely related species, increasing chances for interspecific interactions. When congeneric species frequently occur in the same environments, competition for space, food, or other limiting resources may become more vital in shaping morphological diversity, possibly weakening the extent to which body size tracks temperature alone. Comparative studies on birds show that overlap in thermal environments is closely related to patterns of congeneric co-occurrence and competition, although the relationship between overlap in thermal environments and body-size differential is not always uniform across environments (Srinivasan et al. 2018). Interspecific competition can contribute to habitat and morphological divergence in sympatric areas, as evidenced by closely related songbirds (Sottas et al. 2018). Ecological interactions associated with greater environmental overlap therefore provide a plausible complementary explanation for the association between high thermal similarity and weaker body-size differentiation. However, this mechanism remains speculative in our study and requires direct tests incorporating species co-occurrence, geographic overlap, and interspecific interactions.

The model comparison suggests that thermal similarity captures a more proximal dimension of clinal variation than broad genus-level environmental averages. By summarizing overlap among species-specific temperature distributions, thermal similarity describes how thermal environments are partitioned within a lineage, information that genus-level averages may obscure. Realized thermal conditions are jointly shaped by climate and habitat availability and may therefore approximate the conditions encountered by populations more closely than individual macroclimatic variables (Oboikovitz & Swanson 2022, Conradie et al. 2025). Lower thermal similarity may identify lineages in which congeneric species occupy sufficiently distinct thermal regimes for size-dependent heat exchange to influence divergence consistently. Larger bodies reduce surface area relative to volume and conserve heat under cold conditions, whereas smaller bodies facilitate heat dissipation under warm conditions (Fr hlich et al. 2023, Henry et al. 2023). The support for precipitation points to a complementary pathway. Rainfall influences food availability and growth, and reduced rainfall has been associated with smaller body size in a semi-arid passerine (Gardner et al. 2014). Seasonal precipitation also explained morphological variation better than temperature in an intact Amazonian bird community (Jirinec et al. 2021). Precipitation may additionally modify the water costs of thermoregulation because evaporative cooling becomes critical under high heat loads and scales with body mass (Song & Beissinger 2020). Thermal similarity may thus describe the thermal opportunity for divergence, while precipitation modifies the energetic and hydric conditions under which divergence is expressed.

Beyond thermal similarity alone, our results indicate that ecological context modifies the thermal similarity–slope association. The positive thermal similarity–slope association was supported in dense and semi-open habitats, whereas the estimate for open habitats was more uncertain, and its credible interval included zero. This pattern is consistent with a role for microclimatic buffering. Canopy cover, vertical stratification, and shaded understories buffer forest-dwelling organisms from acute diurnal thermal extremes (Kašpar et al. 2021). By filtering short-term and episodic thermal variation, these structured habitats may allow persistent geographic temperature gradients to exert a clearer influence on body-size divergence (Kim et al. 2022, Strydom et al. 2025). Conversely, birds in open habitats face direct, unbuffered exposure to acute temperature spikes. Under these conditions, fine-scale thermal variability may weaken the correspondence between broad-scale macroclimatic conditions and the temperatures actually experienced by birds (Riddell et al. 2021, Conradie et al. 2025), potentially obscuring the thermal similarity–slope association at the macroecological scale. The smaller number of qualifying genera in the open-habitat subgroup also warrants statistical caution (Gienger et al. 2019, Yang et al. 2022).

In resident birds, a positive association between thermal similarity and body mass–temperature slopes was supported, whereas the corresponding estimate for migratory birds was highly uncertain and its credible interval broadly overlapped zero.

This pattern is consistent with the idea that migratory behavior can act as a behavioral buffer (Mainwaring & Street 2021). Migrant birds may experience weaker or less persistent temperature-mediated selection on body size if they track seasonally favorable environments and limit their exposure to severe winter conditions (Eyres et al. 2020, Cohen & Jetz 2023). Breeding-season temperatures may nevertheless remain ecologically relevant because they characterize the thermal conditions under which breeding occurs. The weaker and more uncertain thermal similarity–slope association in migratory birds could therefore reflect behavioral buffering, variation in thermal environments across the annual cycle, or both. Because our climatic characterization of migratory species was based on breeding ranges, it captures breeding-season thermal conditions but does not represent the full range of thermal environments experienced throughout the annual cycle. Future analyses incorporating climatic conditions across breeding, migration, and non-breeding periods will be important for distinguishing among these explanations.

Additionally, estimating geographical clines for species-poor genera increases uncertainty in genus-level slope estimates. A five-species minimum represents a pragmatic lower threshold here. With four species, genus-level regression has only two residual degrees of freedom, allowing an extreme body mass or thermal position to strongly influence the slope. Mean thermal similarity uses six pairs at *n* = 4 and ten at *n* = 5, improving representation of within-genus thermal structure. This gain is ecologically relevant because congeners differ in habitat use, migration and other strategies that modify body size–temperature relationships (Mainwaring & Street 2021). In a sparsely represented genus, one habitat specialist or geographically peripheral species may therefore cause the estimated cline to reflect lineage composition more strongly than a recurring thermal response. The five-species criterion limits this leverage while retaining enough genera for comparative inference, consistent with the greater stability at the five- and six-species thresholds. Sensitivity analyses using alternative thermal similarity indices and excluding imputed masses further showed that the association was not tied to one overlap metric or body-mass treatment. The subgroup analyses had small sample sizes, particularly for migratory genera (*n* = 7) and open habitats (*n* = 9). These estimates should therefore be interpreted cautiously. While these patterns offer preliminary ecological insights, the high uncertainty (wide credible intervals) in these specific subgroups underscores the need for cautious interpretation, and future studies with broader taxonomic sampling within ecological categories will be essential to verify these context-dependent trends.

Nevertheless, our mechanistic interpretations remain bounded by the intrinsic limitations of global macroecological scaling. First, we relied on gridded macroclimatic data, which inherently smooth over the microscale thermal realities that small, behaviorally plastic birds actually experience (Riddell et al. 2021, Conradie et al. 2025). Furthermore, birds employ a range of physiological and morphological mechanisms that can buffer thermal stress, including plumage-mediated insulation, facultative torpor, and metabolic flexibility, none of which were incorporated in the present analysis (McKechnie et al. 2023). These unmeasured physiological and behavioral traits may represent additional sources of variation in genus-level body mass–temperature slopes. Resolving this will heavily depend on the future integration of high-resolution microclimate grids with massive, multi-trait functional databases.

Overall, our findings provide a new macroecological perspective on variation in the strength of Bergmann’s rule across avian genera. By linking variation in thermal similarity to heterogeneity in Bergmannian clinal strength, our study provides an ecological framework that may help explain why closely related bird lineages differ in their morphological responses to climatic gradients. This framework further suggests that the expression of temperature-associated body-size divergence is related to the degree of thermal differentiation among congeneric species, with habitat structure and migratory behavior potentially modifying this association. Future studies integrating full-annual-cycle climatic exposure, high-resolution microclimatic data, and long-term morphological records will be important for directly testing these proposed mechanisms and determining whether thermal similarity can improve forecasts of avian morphological responses under ongoing climate change. Extending this framework to other endothermic clades would further clarify how generally variation in thermal similarity contributes to variation in ecogeographical body-size patterns.

## Data Availability Statement

The analysis-ready derived datasets, metadata, and custom R and Python code required to reproduce the analyses, sensitivity analyses, model comparisons, tables, and figures are available during peer review via an anonymized private Figshare link (https://figshare.com/s/d0a3935904e90a4e23dd). These materials will be made publicly available on Figshare upon publication under a persistent DOI. The species-level source dataset was obtained from He et al. (2023) and its Dryad archive. Original third-party data, including the BirdLife International and NatureServe avian range polygons, are not redistributed and remain subject to the access and licensing terms of their providers; complete source citations and access information are provided in the Materials and Methods.

## Supporting Information

The Supporting information associated with this article is available with the online version.

